# Microtubule acetylation but not detyrosination promotes focal adhesion dynamics and cell migration

**DOI:** 10.1101/425421

**Authors:** Bertille Bance, Shailaja Seetharaman, Cécile Leduc, Batiste Boëda, Sandrine Etienne-Manneville

## Abstract

Microtubules play a crucial role in mesenchymal migration by controlling cell polarity and the turnover of cell adhesive structures on the extracellular matrix. The polarized functions of microtubules imply that microtubules are locally regulated. Here, we investigated the regulation and role of two major tubulin post-translational modifications, acetylation and detyrosination, which have been associated with stable microtubules. Using primary astrocytes in a wound healing assay, we show that these tubulin modifications are independently regulated during cell polarization and differently affect cell migration. In contrast to microtubule detyrosination, αTAT1-mediated microtubule acetylation increases in the vicinity of focal adhesions and promotes cell migration. We further demonstrate that αTAT1 increases focal adhesion turnover by promoting Rab6-positive vesicle fusion at focal adhesions. Our results highlight the specificity of microtubule post-translational modifications and bring new insight into the regulatory functions of tubulin acetylation.

## Introduction

Cell migration relies on the polarization and coordinated regulation of numerous cell structures, including cytoskeletal elements and adhesive structures (Llense and Etienne-Manneville, 2015, Wolfenson et al., 2009, Gardel et al., 2010, Parsons et al., 2010). While actin plays a crucial role in the generation of forces that promote cell protrusion and cell net displacement, microtubules participate in cell front-to-rear polarization (Elric and Etienne-Manneville, 2014, Etienne-Manneville, 2013) and focal adhesion dynamics (Stehbens and Wittmann, 2012, Etienne-Manneville, 2013) to promote astrocyte or fibroblast migration and endothelial cell invasion (Etienne-Manneville, 2004, Etienne-Manneville and Hall, 2001, Bouchet et al., 2016, Bouchet and Akhmanova, 2017). At the cell front, microtubules are captured in the vicinity of nascent adhesions and contribute to the polarized delivery of integrins towards these sites (Bretscher and Aguado-Velasco, 1998). Microtubules can also target focal adhesions (Kaverina et al., 1998) and induce their disassembly through the recruitment of the endocytic machinery (Kaverina et al., 1999, Kaverina et al., 2002, Palazzo et al., 2004, Ezratty et al., 2009, Stehbens et al., 2014). The polarized organization of integrin-based structures implies that the functions of microtubules must be controlled in a polarized manner.

Microtubule organization, dynamics and functions are regulated by several mechanisms including their interaction with MAPs and post-translational modifications of α- and β-tubulin (Janke, 2014, Song and Brady, 2015, Etienne-Manneville, 2010, Strzyz, 2016) (Aillaud et al., 2016) that can affect microtubule dynamics or its association with protein partners (Janke, 2014, Raunser and Gatsogiannis, 2015, Yu et al., 2015, Song and Brady, 2015). Lysine 40 acetylation and carboxy-terminal detyrosination of alpha tubulin are both classically used as markers of stabilized microtubules, which are relatively long-lived and resistant. Both detyrosinated and acetylated microtubules accumulate between the centrosome and the leading edge of migrating fibroblasts (Gundersen and Bulinski, 1988b, Montagnac et al., 2013, Castro-Castro et al., 2012). However, the specific functions of detyrosination and acetylation of microtubules are not well understood (Gadadhar et al., 2017, Song and Brady, 2015). Alpha-tubulin acetylation has been shown to facilitate fibroblast and neuronal microtubule-dependent motility (Hubbert et al., 2002, Creppe et al., 2009) but increased acetylation of microtubules can also reduce cell migration (Tran et al., 2007). Using primary rat astrocytes, whose migration is highly dependent on microtubules, we analyzed and compared the regulation and roles of microtubule acetylation and detyrosination. We found that these two tubulin modifications are independently regulated and that tubulin acetylation specifically impacts cell migration and increases focal adhesion turnover by promoting Rab6A vesicle fusion at focal adhesions.

## Results and Discussion

### Microtubule acetylation but not detyrosination is up-regulated during cell polarization and promotes cell migration

To determine how acetylation and detyrosination were regulated during migration we quantified acetylated and detyrosinated microtubules before and during wound-induced cell polarization and migration using antibodies specifically directed against anti-detyrosinated tubulin and K40 acetylated tubulin. In confluent non migrating astrocytes, acetylated and detyrosinated microtubules are concentrated in the perinuclear region (Fig. 1A, 1B). Wound-induced polarization of astrocytes induced an increase in the proportion of acetylated tubulin and in parallel, a decrease in the proportion of detyrosinated tubulin (Fig. 1A, 1B and S1A). The changes were maximum 8h after wounding when the cells are fully polarized and migrate at a steady state (Fig. 1A, 1B). The decrease in microtubule detyrosination differs from what was observed during wound-induced fibroblast migration (Gundersen and Bulinski, 1988a). However, fibroblast migration requires cell stimulation by serum or LPA and microtubule detyrosination was shown to increase in response to LPA-induced RhoA activation (Cook et al., 1998). During astrocyte wound-healing assay, serum or LPA are not added and polarization and migration result solely from the wound of the cell monolayer (Etienne-Manneville, 2006). Using structure illumination based super-resolution microscopy to distinguish acetylation and detyrosination on single microtubules, we observed that tubulin acetylation and detyrosination were frequently found on the same microtubules. However, they did not always colocalize, often decorating distinct microtubule regions (Fig. 1C).

**FIGURE 1:**
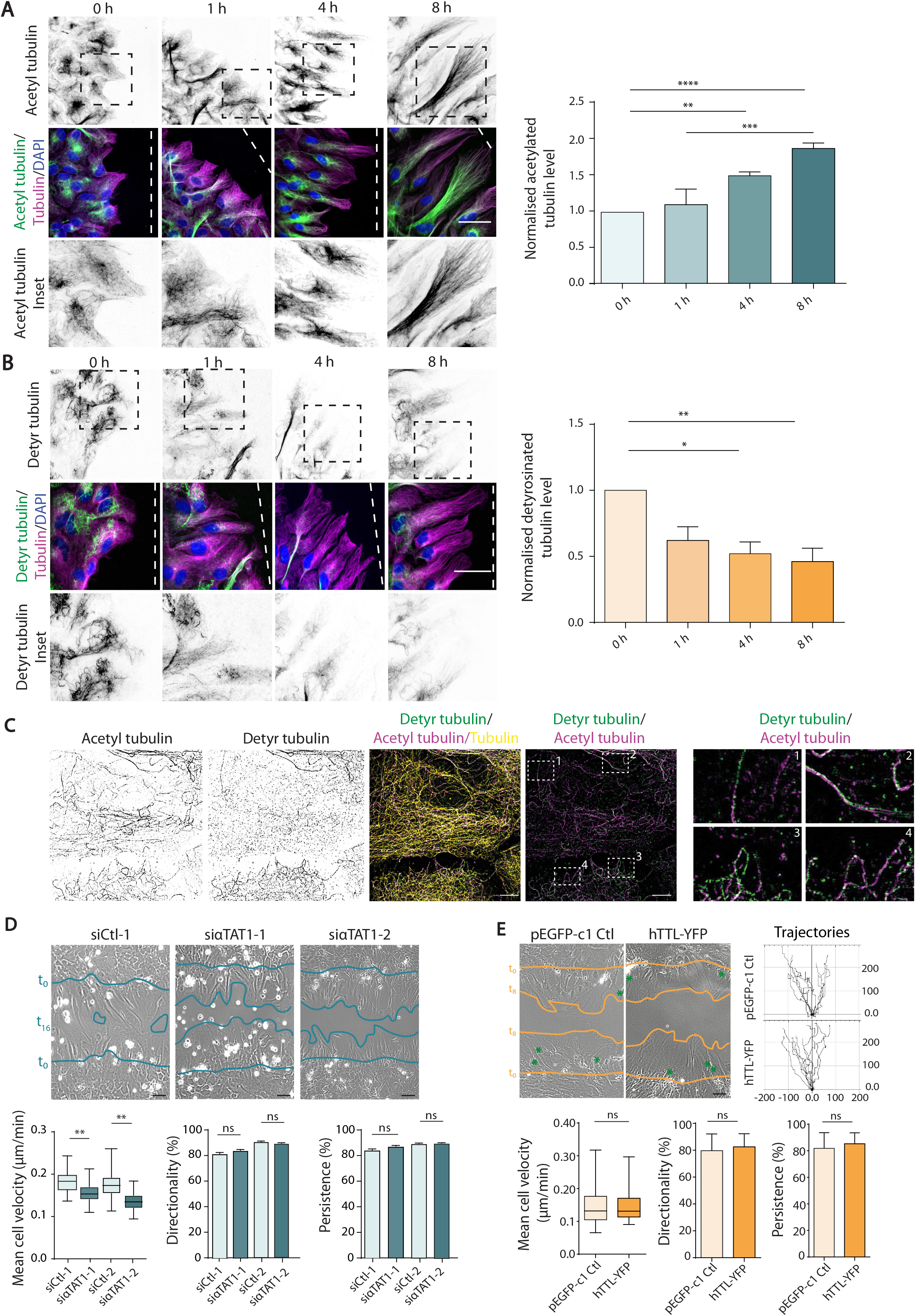
Microtubule acetylation but not detyrosination is increased during wound-induced migration and promotes cell migration while microtubule detyrosination inhibits cell migration. (A, B) Migrating astrocytes fixed at indicated time points following wounding were stained with antibodies against α-tubulin, acetylated tubulin (A) or detyrosinated tubulin (B). Dashed lines indicate the wound orientation. Scale bar: 50 µm. Graph represents the mean normalized acetyl tubulin intensities for each condition ± SEM; for acetyl levels, n ≥ 286 cells/condition from 3 independent experiments; for detyrosinated tubulin levels, n ≥ 128 cells/condition from 3 independent experiments; one-way ANOVA, Tukey’s multiple’s comparison’s test. (C) Super-resolution images showing astrocytes in a monolayer, stained for detyrosinated tubulin (green), acetylated tubulin (magenta) and α-tubulin (yellow). White dashed boxes show regions of the zooms (on the right). Scale bar: 10 µm. (D) Images of migrating astrocytes, transfected with indicated siRNAs at 16 h, after wounding. Blue lines marked t_0_ indicate the initial wound edge and t_16_ indicates wound gap after 16 h. Scale bar: 100 µm. The box-and-whisker plot show the cell velocity (µm/min), directionality (%) and persistence (%) of migrating astrocytes (as described in the material and methods section); n ≥ 70 cells from 3 independent experiments); oneway ANOVA. (E) Images of migrating astrocytes expressing pEGFP-c1 or hTTL-YFP, 8 h post wounding. Blue lines marked t_0_ indicate the initial wound edge and t_8_ indicates wound gap after 8 h. Scale bar: 100 µm. Trajectories of 10 cells each from pEGFP-c1 and hTTL-YFP are shown (right). Graph shows mean cell velocity ± SEM (µm/min), directionality (%) and persistence (%) of migrating astrocytes; n ≥ 70 cells from 3 independent experiments; one-way ANOVA. The box-and-whisker plots show median, first and third quartiles (boxes), whiskers extending to the furthest observations. ns – not significant, * p < 0.05, ** p < 0.01, *** p < 0.001, **** p< 0.0001.

We then determined the role of microtubule acetylation on cell migration. αTAT1 (also known as Mec-17 in C. elegans (Akella et al., 2010)) is the major tubulin acetyltransferase in mammals (Kalebic et al., 2013b). We used two distinct sets of 2 siRNAs directed against different exons of the αTAT1 acetylase. αTAT1 depletion strongly decreased microtubule acetylation (Fig. S1B) without modifying detyrosinated tubulin level (Fig. S1D, S1E). Cell treatment with tubacin, a specific HDAC6 inhibitor (Haggarty et al., 2003) which dramatically increases microtubule acetylation did not affect detyrosination (Fig. S1D, S1E). siRNA-mediated depletion of αTAT1, decreased the cell migration speed but did not affect the direction or the persistence of migration (Fig. 1D, movie 1). The second set of siRNA directed against αTAT1 had a similar effect on migration as compared to a non-relevant siRNA (Fig 1D, movie 1). The polarized shape of wound edge cells was not perturbed by the reduction of acetylated microtubules (Fig. S2A, movie 1).

Detyrosination is the elimination of the C-terminal tyrosine residue from α-tubulin by tubulin carboxypeptidase TCP (Aillaud et al., 2017). The tubulin tyrosine ligase (TTL) catalyzes the reverse reaction on soluble tubulin. To assess the specific role of detyrosination in comparison with acetylation, we decreased detyrosinated tubulin by overexpressing the tubulin tyrosine ligase hTTL (Fig. S1C). Tracking of hTTL-YFP expressing cells showed that tubulin detyrosination does not perturb the cell migration speed, direction nor persistence (Fig 1E, movie 2). Altogether, our results indicate that microtubule detyrosination and acetylation are independently regulated during cell polarization and have distinct roles during cell migration.

### Microtubule acetylation but not detyrosination affects focal adhesions

Since microtubules have been shown to be crucial for the control of cell adhesion, we used paxillin staining to investigate the impact of microtubule acetylation and detyrosination on focal adhesions in migrating cells. Depletion of αTAT1 increased the number of focal adhesions (+35% compared to siRNA control nucleofected cells) without modifying the focal adhesion size (Fig. 2A, 2B). Moreover, inhibition of microtubule acetylation altered the distribution of focal adhesions. In control cells, focal adhesions are restricted to the front of the protrusion of wound edge cells, whereas in αTAT1-depleted cells, focal adhesions visible throughout the basal surface, resulting in an increased mean distance between focal adhesions and the cell leading edge (Fig. 2C). Expression of a siRNA-resistant αTAT1 construct rescued the number and distribution of focal adhesions whereas a siRNA-resistant construct encoding a catalytically inactive αTAT1 (GFPα-TAT1-D157N) did not rescue the phenotype (Fig. 2A, 2C). In contrast to acetylation, reducing microtubule detyrosination did not affect focal adhesion number, size or distribution (Fig. 2B, 2D), showing that microtubule acetylation, but not detyrosination, in specific, affects focal adhesion dynamics.

**FIGURE 2:**
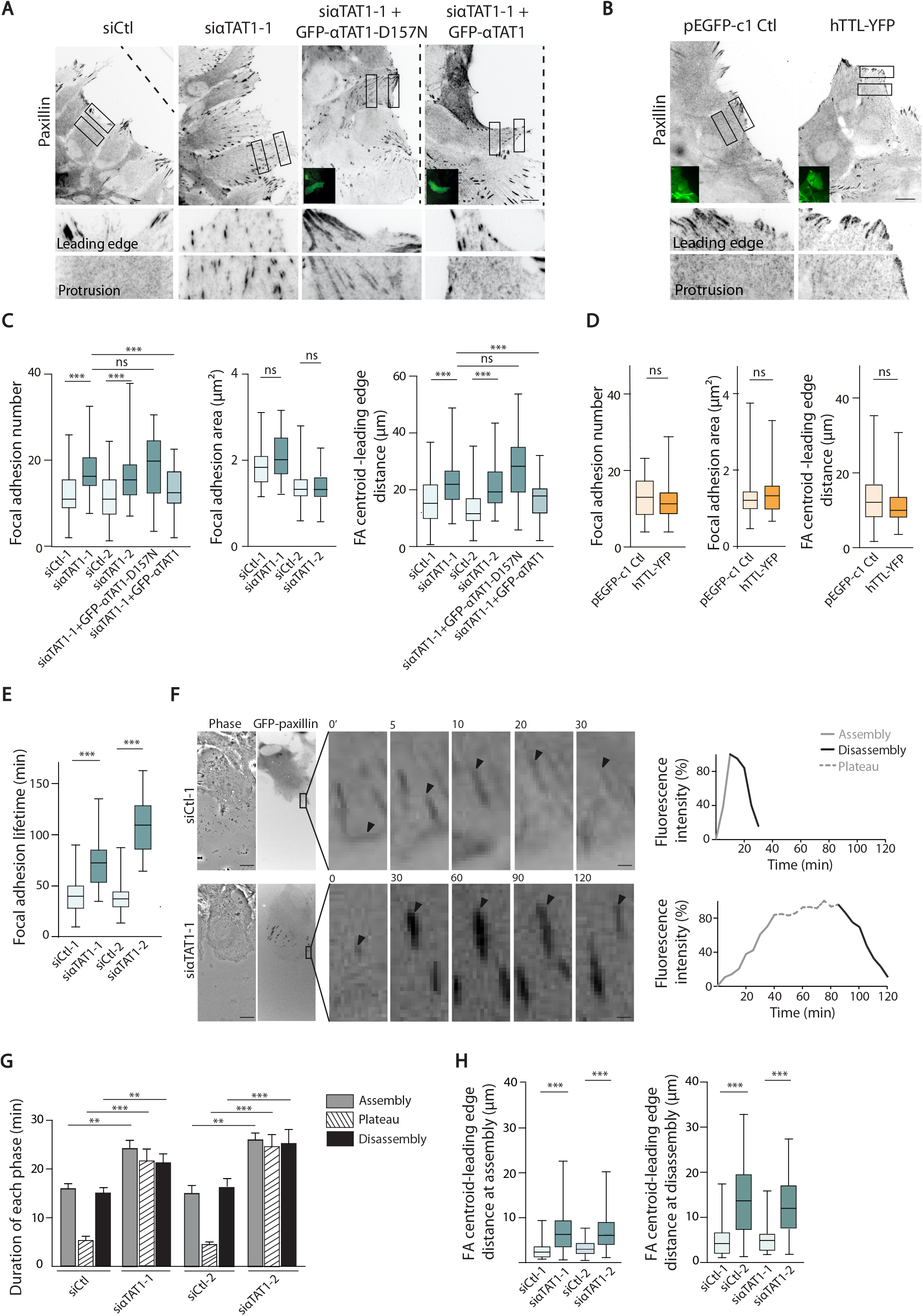
Microtubule acetylation but not detyrosination controls focal adhesions. (A, B) Epifluorescence images of paxillin immunostaining (inverted contrast) in migrating (8 h after wounding) astrocytes nucleofected with the indicated siRNAs and/or plasmids (the insets show GFP expression). Dashed lines indicate the wound orientation. Scale bars: 10 µm. The bottom panels correspond to the zooms of the boxed regions located at the leading edge or further back within the protrusions. Scale bar: 1 μm. (C, D). Graphs show the mean FA number, FA area (µm^2^) and the distance between FAs and leading edge (µm) in migrating astrocytes transfected with indicated siRNAs and plasmids (n > 3, 100 cells per conditions/per experiment). Statistical significance was determined using a one-way ANOVA. (E) Box-and-whisker plot showing the focal adhesion lifetime in migrating astrocytes transfected with GFP-Paxillin and indicated siRNAs. (n = 3; 10 focal adhesions per cell; 10 cells per condition). (F) The two left panels show the phase contrast and fluorescence (inverted contrast) images of migrating astrocytes expressing GFP-Paxillin (Movie 2; total time, 4 hours). In each case, five images taken from the time lapse movies (real time (in min) are indicated in the top left corners) show the dynamics of focal adhesions located in the region (marked with a black box) of the corresponding GFP-paxillin expressing cells. Scale bar: 5 µm (0.5 µm in the inset). The graphs on the right show the fluorescence intensity over time of one focal adhesion (indicated by a black arrowhead) during its phase of assembly, plateau and disassembly phase. The fluorescence intensity is given as a percentage of the maximum intensity. (G) Boxand-whisker plot showing distance between focal adhesion centroid and leading edge at the different phases of assembly, plateau and disassembly of GFP-paxillin in migration astrocytes transfected with indicated siRNAs. Histograms show mean ± SEM (n = 3, 5 focal adhesion per cell, 10 cells per condition). (H) Graph indicates distance between the focal adhesion centroid and the leading edge (µm) of migrating astrocytes transfected with siRNA (n > 3, 100 cells per conditions per experiment). The box-and-whisker plot show median, first and third quartiles (boxes), whiskers extending to the furthest observations. Histograms show mean ± SEM; Student’s t test. ns – not significant (p>0.05), ** p < 0.01, *** p < 0.001.

We then used paxillin-GFP expressing astrocytes to analyse focal adhesion dynamics. αTAT1-depletion increased the focal adhesion lifetime as compared to control cells (Fig. 2E, movie 3). In control astrocytes, focal adhesion turnover is composed of two steps: assembly and disassembly. During the phases of assembly and disassembly, focal adhesion area increases or decreases over time respectively (Gardel et al., 2010) (Fig. 2F). Decreased microtubule acetylation was associated with longer assembly and disassembly phases separated by a phase during which the size of focal adhesions plateaued (Fig. 2F, 2G). Moreover, in agreement with the wider distribution of focal adhesions in αTAT1-depleted cells, the distance to the leading edge of assembling and disassembling focal adhesions was increased in αTAT1-depleted cells (Fig. 2H). Altogether, these results show that the tubulin acetylase αTAT1 specifically controls focal adhesion number, distribution and dynamics.

### Acetylated microtubules and αTAT1 localize at focal adhesions

The impact of αTAT1 on focal adhesion dynamics led us to investigate the localization of αTAT1 and acetylated microtubules. Cell wounding initially increased microtubule acetylation in the perinuclear region (Fig. 3A, 3B). Between 4h and 8h after wounding, acetylated microtubules further accumulated towards the cell front (Fig. 3A, 3B). Stretches of acetylated tubulin were concentrated on microtubules in proximity to focal adhesions at the front of migrating cells while they appear less abundant in region without focal adhesions (Fig. 3C).

**FIGURE 3:**
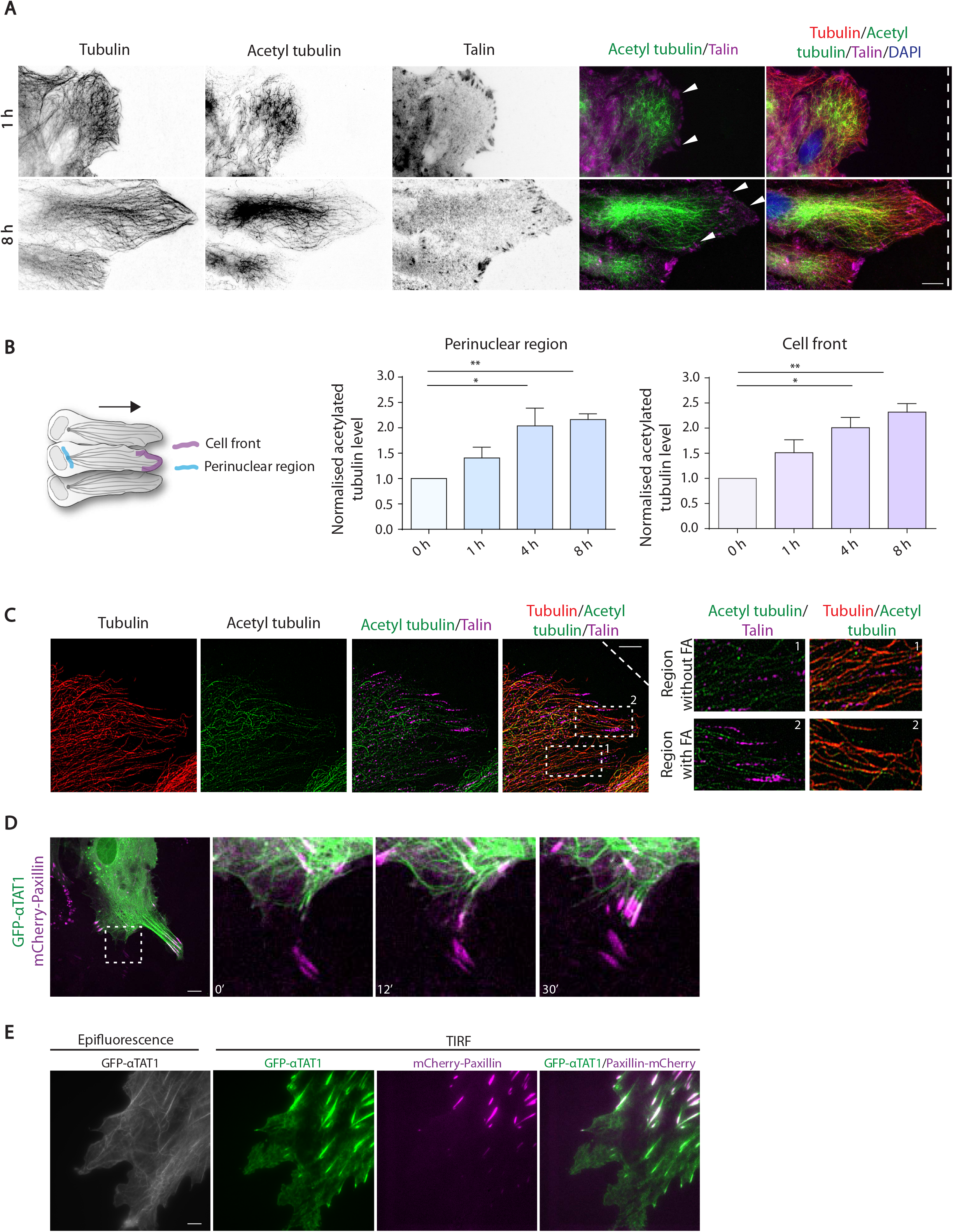
Acetylated microtubules and αTAT1 acetyltransferase localize at focal adhesions. (A) Epifluorescence images of migrating astrocytes at different time points after wounding, immunostained with α-tubulin (inverted contrast and red), acetylatedtubulin (inverted contrast and green) and talin (inverted contrast and magenta). Dashed lines indicate wound orientation and white arrows indicate regions of interest near focal adhesions. Scale bars: 10 µm. (B) Graphs showing mean ± SEM of the normalized ratios between acetylated tubulin and total tubulin at the cell front (purple) or perinuclear region (blue). Quantifications of intensities in the different regions are as shown in the left schematic, at different time points following wounding. For perinuclear acetyl levels, n ≥ 62 cells from 3 independent experiments; for acetyl levels at cell front, n ≥ 167 cells from 3 independent experiments. (C) Super-resolution images of α-tubulin (red), acetylated-tubulin (green) and talin (magenta) immunostaining of a migrating astrocyte 8h after wounding. Scale bar: 10 µm. The higher-magnification images of regions indicated by white dashed boxes are shown on the right. (D) Fluorescence images of a GFP-αTAT1 (green) and Paxillin-mCherry (magenta) expressing astrocyte 6 h after wounding (see movie 5). The higher-magnification images of region indicated by a white dashed box at different time points are shown at the right. Scale bar: 5 µm (1 µm in the zoom). (E) Epifluorescence (left) and TIRF images of a GFP-αTAT1 (grey or green) and Paxillin-mCherry (magenta) expressing astrocyte. Scale bar: 5 µm.

Since an anti-αTAT1 antibody did not allow us to visualize the endogenous protein in fixed cells, we used GFP-αTAT1 expression to determine the localization of αTAT1. Expression of GFP-αTAT1 alone did not alter focal adhesion number nor distribution (Fig. S2B) but it rescued focal adhesions numbers in αTAT1-depleted cells (Fig. 2A, 2B), suggesting that it behaves like the endogenous protein. GFPα-TAT1 colocalized with microtubules (Fig. 3D and movie 4). Moreover, TIRF microscopy revealed an accumulation of GFP-αTAT1 in paxillin-mCherry positive focal adhesions (Fig. 3E, movie 5). These observations suggest that αTAT1 localizes at focal adhesions and can locally promote acetylation of microtubules. The regulation of αTAT1 localization and/or activity during migration remains to be elucidated but may involve integrin signaling (Palazzo et al., 2004) or αTAT1 interaction with clathrin-coated pits localized around focal adhesions (Montagnac et al., 2013).

### αTAT1 controls Rab6 vesicle fusion at focal adhesions

The targeting of focal adhesions by microtubules have been shown to control focal adhesion dynamics through vesicular traffic (Ezratty et al., 2009, Ezratty et al., 2005, Shafaq-Zadah et al., 2016), suggesting that microtubule acetylation effect on focal adhesion dynamics may involve the regulation vesicular traffic. The small GTPase Rab6 decorates most post-Golgi carriers (Fourriere et al., 2018). Moreover Rab6 and Rab8 have been involved in vesicular traffic to and from focal adhesions (Stehbens et al., 2014, Grigoriev et al., 2007). Rab6A-positive vesicles transported along microtubules have been shown to fuse at cortical focal adhesion associated sites to control focal adhesion dynamics (Grigoriev et al., 2011). We thus expressed GFP-Rab6A in primary astrocytes to assess the impact of αTAT1 on Rab6A-positive vesicles. GFP-Rab6A associated with the Golgi complex and cytoplasmic vesicles as described in other cell types (Goud et al., 1994). Most of Rab6A-positive vesicles appeared to move from the Golgi apparatus to the cell periphery (movie 6) and may correspond to constitutive secretion carriers (Grigoriev et al., 2007, Grigoriev et al., 2011). Using Paxillin-mCherry expressing cells, we observed that GFP-Rab6A positive vesicles were transported towards focal adhesions where the fluorescence disappeared indicating vesicle fusion with the plasma membrane (Grigoriev et al., 2011)(Fig. 4A, movie 6). Kymograph analysis showed that in control cells GFP-Rab6A vesicles underwent rapid docking followed by fusion. In contrast, GFP-Rab6A vesicles in αTAT1-depleted cells reached focal adhesions but the number of fusion events was strongly decreased (Fig. 4A, 4B). Furthermore, when fusion occurred, the time between immobilization and actual fusion with plasma membrane was strongly increased in the case of αTAT1-depleted cells (Fig. 4C, persistence time). To determine the role of Rab6-dependent traffic on focal adhesion dynamics, we used specific siRNA to downregulate of Rab6A expression (Fig. 4D). Rab6A depletion increased the number of focal adhesions and altered their distribution in a manner similar to that observed in αTAT1-depleted cells (Fig. 4E, 4F), suggesting a functional relationship between αTAT1-mediated microtubule acetylation and Rab6A-dependent vesicular traffic. Rab6A controls the transport and fusion of secretory carriers (Grigoriev et al., 2007, Grigoriev et al., 2011) and we can hypothesise that it contributes to the trafficking of essential components towards the forming focal adhesions, which may explain the slower assembly of focal adhesions in the absence of αTAT1. Rab6A has also been involved in the retrograde transport of integrins during cell migration (Shafaq-Zadah et al., 2016), suggesting that microtubule acetylation may also affect the focal adhesion dynamics by controlling integrin recycling. Altogether, these observations strongly suggest that αTAT1-mediated microtubule acetylation facilitate Rab6A-positive vesicle fusion to promote focal adhesion dynamics.

**FIGURE 4:**
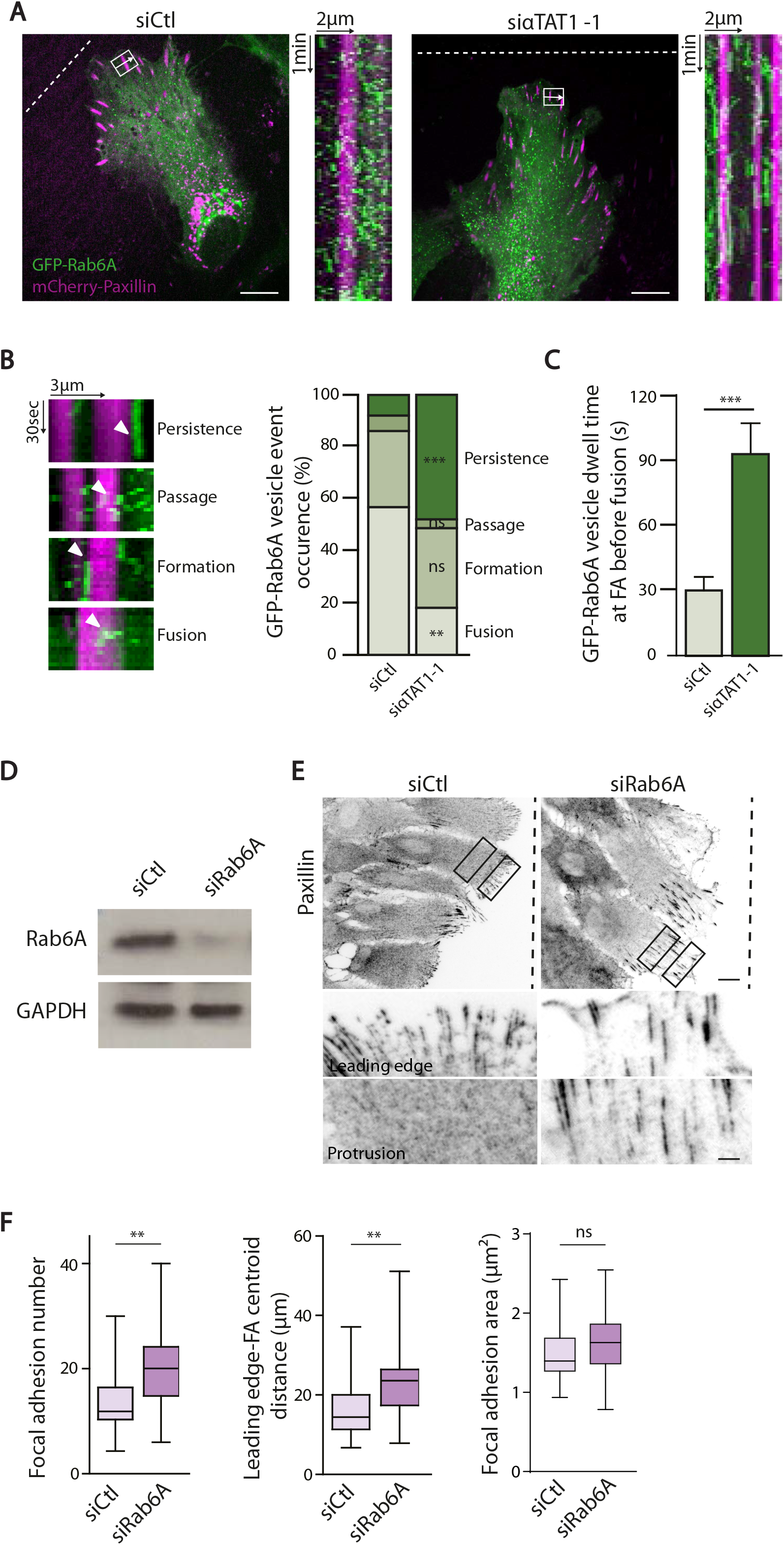
αTAT1 controls Rab6A-positive vesicle fusion at focal adhesions. (A) Fluorescence images of migrating astrocytes nucleofected with indicated siRNAs, GFP-Rab6A (green) and Paxillin-mCherry (magenta) (movie 6, total time: 10 min). Kymographs (to the right of each image) show the fluorescence intensity profile along the white arrow shown in the white box in the corresponding image over time. Scale bar: 10 µm. (B) Graph showing the percentage of the different behaviors (persistence, passage, formation, fusion) of GFP-Rab6A vesicles observed on the kymograph. (C) Graph showing dwell time of GFP-Rab6A vesicles at focal adhesions before disappearance. n > 50 FAs from three independent experiments (6 cells per experiment and > 50 GFP-Rab6A vesicles trajectories analysed). (D) Western blot analysis with indicated antibodies of total protein lysates of astrocytes nucleofected with the indicated siRNAs or plasmids. The result shown is representative of 3 independent experiments. (E) Epifluorescence images of paxillin (inverted contrast) in migrating astrocytes (8 h after wounding) nucleofected with the indicated siRNAs. Dashed lines indicate wound orientation. Scale bar: 10 µm. The bottom images show zooms of regions indicated by the dashed boxes. Scale bar: 2 µm. (F) Graphs showing the number, area and distance to the leading edge (µm) of FAs in migrating astrocytes transfected with indicated siRNAs. The box-and-whisker plot show median, first and third quartiles (boxes), whiskers extending to the furthest observations. n > 50 cells per condition from 3 independent experiments, 5 to 10 focal adhesions per cell; one-way ANOVA.

Our results highlight the specific regulation and role of microtubule acetylation and detyrosination during cell migration. The fact that detyrosination and acetylation during astrocyte migration are differently regulated and can be found on distinct microtubule regions strongly suggest that these two modifications often considered as equivalent markers of stable microtubules are, in fact, very unique. Moreover, microtubule acetylation, but not detyrosination, is associated with faster focal adhesion turnover and cell migration. So far, αTAT1 is the only acetyl transferase known to induce tubulin acetylation of lysine 40 and no other substrates of αTAT1 are known. The effects of αTAT1 depletion were systematically correlated with decreased acetylation levels, which lead us to conclude that the function of αTAT1 is due to microtubule acetylation. However, we cannot completely exclude the possibility that αTAT1 also acts through an alternative mechanism. αTAT1 has been shown to destabilize microtubules independently of its acetyltransferase activity (Kalebic et al., 2013a) and its interaction with proteins such as doublecortin or cortactin may be involved (Kim et al., 2013, Castro-Castro et al., 2012). Nevertheless, the effects of αTAT1 depletion of focal adhesion turnover were rescued by expression of αTAT1-WT but not by the catalytically inactive mutant, confirming the role of acetylation in αTAT1 functions.

Altogether, our results demonstrate the specificity of microtubule acetylation and detyrosination, two post-translational modifications identically used at markers of stable, long lived microtubules. We also show here that microtubule acetylation plays a crucial role in the control of vesicular traffic, focal adhesion dynamics and ultimately control the cell migration speed.

## Materials and Methods

### Cell culture

Primary astrocytes were obtained from E17 rat embryos (Etienne-Manneville, 2006). They were grown in 1g/L glucose DMEM supplemented with 10% FBS (Invitrogen, Carlsbad, CA), 1% penicillin-streptomycin (Gibco) and 1% Amphotericin B (Gibco) at 5% CO_2_ and 37°C.

### Transfection

Astrocytes were transfected with Lonza glial transfection solution and electroporated with a Nucleofector machine (Lonza). Cells were then plated on appropriate supports previously coated with poly-L-Ornithine (Sigma). Experiments are carried out 3 or 4 days post-transfection and comparable protein silencing was observed. siRNAs were used at 1 nmol for astrocytes. siRNA sequences used were: Luciferase (control): UAAGGCUAUGAAGAGAUAC. αTAT1 rat (siαTAT1-1): 5’-ACCGACACGUUAUUUAUGU-3’ and 5’-UUCGAAACCUGCAGGAACG-3’. αTAT1 rat (siαTAT1-2): 5’-UAAUGGAUGUACUCAUUCA-3’ and 5’-UCAUGACUAUUGUAGAUGA-3’. Rab6A rat: 5’-GCA ACA AUU GGC AUU GACUUC UUA U-3’ and 5’-CAA ACA AUU CCA GCA UAC UCG UGA U-3’.

DNA plasmids used were: pEGFP (Clontech), pEGFP-paxillin (from E.M. Vallés, Institut Curie), pPaxillin-mCherry cloned from pPaxillin psmOrange with BamHI and NotI, hTTL-GFP (human) (from Carsten Janke, Institut Curie, Orsay), EGFP-Rab6A (human) (from Bruno Goud, Institut Curie, Paris), EGFP-αTAT1 and GFP-αTAT1-D157N (from Philippe Chavrier, Institut Curie, Paris).

### 2D wound healing assay

Cells were plated on appropriate supports (dishes, plates, coverslips or glass-bottom MatTek). Cells were allowed to grow to confluence and fresh medium was added the day prior to the experiment. The cell monolayer was then scratched with a p200 pipette tip to induce migration.

### Immunofluorescence

Cells migrating for 8h (unless otherwise stated) were fixed with cold methanol for 3-5 min at −20°C or 4% warm PFA for 10 min and permeabilised for 10 min with Triton 0.1%. Coverslips were blocked for 30 min with 5% BSA in PBS. The same solution was used for primary and secondary antibody incubation for 1 h. Nuclei were stained with DAPI or Hoechst and were mounted with Prolong Gold with DAPI (Life Technologies). Epifluorescence images were acquired with a Leica DM6000 microscope equipped with 40X 1.25 NA or 63X 1.4 NA objectives and recorded on a CCD camera with a Leica software. Super-resolution 3D-SIM images were acquired with a Zeiss LSM780 ELYRA with 63X 1.4 NA or 100X 1.46 NA objectives and recorded on an EMCCD camera Andor Ixon 887 1K with Zen software.

Antibodies used in this study are anti-acetylated tubulin (clone 6-11B-1, Sigma), Poly-Glu tubulin (ValBiotech AbC0101), α-tubulin (Biorad MCA77G), GFP-FITC (Abcam), Paxillin (BD transduction), Talin (Sigma T3287). For live experiment, microtubules were fluorescently labelled with Sir-Tub (Cytoskeleton, 1/2500 of a 50µM stock) 1h prior image acquisition.

### Live imaging

For phase contrast wound healing assays, cells in 12 well plates were wounded with a p200 pipette tip and directly placed in the microscope with the addition of HEPES and paraffin to eliminate medium evaporation. Image acquisition was launched about 30 min after wounding. Movies were acquired with a Zeiss Axiovert 200M equipped with a thermostatic humid chamber with 5% CO_2_ and 37°C. Images were acquired every 15 min for 24h, with dry objective 10X 0.45 NA and an EMCCD or sCMOS pco edge camera.

For fluorescence live imaging, cells on appropriate dishes (MatTek) were wounded and allowed to migrate 4h before image acquisition. HEPES and antioxidants were added to the medium before acquisition. Epifluorescence experiments investigating paxillin turnover were performed with a Nikon BioStation IM-Q (40X objective) at a rate of 1 frame/5 min for 3-5h with 5% CO_2_ and 37°C. Confocal experiments were performed on a Perkin-Elmer spinning disk confocal microscope equipped with an EMCCD camera and a 63X 1.4 NA objective with 5% CO_2_ and 37°C. Super-resolution experiments were acquired with the Zeiss LSM780 ELYRA. Images were processed for Structured Illumination with Zen software.

### Electrophoresis and Western blot

Cells lysates were obtained with Laemmli buffer composed of 60 mM Tris-HCl pH6.8, 10% glycerol, 2% SDS and 50 mM DTT with the addition of anti-protease (cOmplete cocktail, Roche 11 873 588 001), 1 mM glycerol phosphate, 1 mM sodium orthovandate and 1 mM sodium fluoride. Samples were boiled 5 min at 95°C before loading on polyacrylamide gels. Transfer occurred at 100V for 1 h on nitrocellulose membranes. Membranes were blotted with TBST (0.2% Tween) and 5% milk and incubated for 1 h with the primary antibody, followed by 1 h with HRP-conjugated secondary antibody. Bands were revealed with ECL chemoluminescent substrate (Pierce, Thermo Scientific or Biorad).

Antibodies used were: anti-α-tubulin (Biorad MCA77G), GAPDH (Chemicon International MAB374), anti-acetylated tubulin (clone 6-11B-1, Sigma), Rab6A (Santa Cruz). Secondary antibodies were all from Jackson ImmunoResearch.

### Quantifications of images and movies

Analysis of focal adhesion number, area and distribution was performed with a custom-designed macro in Fiji. Briefly, for each image focal adhesion contours were determined by thresholding the paxillin fluorescence channel and using the Analyze Particles plugin and a minimum size of 0.5 µm².

To quantify focal adhesion dynamics, integrated fluorescent intensity of single focal adhesion was analysed over time. The turnover was defined as the time elapsed between the appearance (first frame) and the disappearance (last frame). Assembly was defined as the time between appearance and the peak or plateau of fluorescence intensity. The plateau was defined as the period during which the integrated fluorescence of focal adhesion was high and constant, and disassembly was defined as the time between the beginning of integrated fluorescence intensity decrease and disappearance of the focal adhesion.

Normalised mean intensity levels of acetyl and detyr tubulin for immunofluorescence images are calculated as follows:

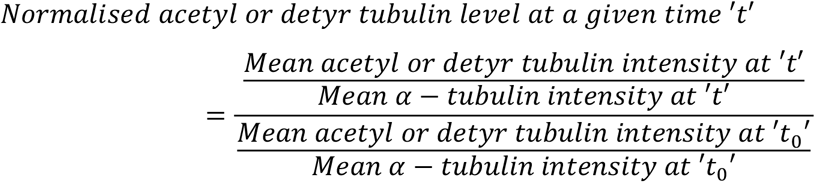

Mean velocity (*v*), persistance (*p*) and directionality (*d*) of cell migration are calculated as follows: for a given (*x*, *y*) coordinate of leading cell nucleus,

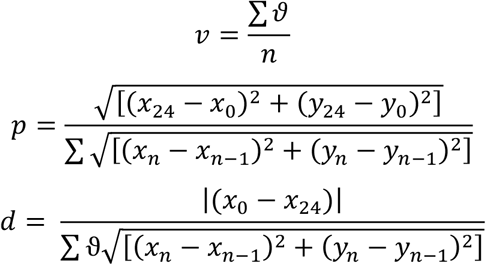

where *n* is the number of time points acquired and *ϑ* is cell velocity.

Mean acetylated tubulin levels at perinuclear or cell front is calculated by drawing a line (width 50) around the specified regions and as shown in Fig. 3B.

For GFP-Rab6A vesicle analysis (Fig 4), ROI of approximately 10-20 µm^2^ around single focal adhesions were drawn using Metamorph. Areas were selected in the Paxillin-mCherry channel independent of the GFP-Rab6A channel to not bias region selection. The image sequence was then cropped closely around FAs as indicated by the box in Fig 4A. A sequence of x-t kymographs was generated with Metamorph. The resulting image sequence is shown in Fig 4A. GFP-Rab6A vesicle events were separated in 4 categories. Fusion corresponds to the disappearance of the GFP-Rab6A vesicle which stopped moving close to the FAs. Persistence corresponds to GFP-Rab6A vesicles that do not immediately disappears but remain close to the FAs before disappearing. Note, that persistence of vesicles in the proximity of FAs is only observed in αTAT1-depleted cells. Formation corresponds to the appearance of the vesicle close to the FAs and passage corresponds to a GFP-Rab6A vesicle moving across FAs. GFP-Rab6A vesicle dwell time was defined as the persistence time of a vesicle before fusion and was measured as the length of vertical GFP-Rab6A tracks in the kymographs. GFP-Rab6A vesicle behavior and dwell-time were quantified. n > 50 FAs from three independent experiments (6 cells per experiment and > 50 GFP-Rab6A vesicles trajectories analysed). Statistical differences were determined using Student’s t-test for GFP-Rab6A vesicles dwell time and a χ^2 by^ contingency for the percentage of vesicle events.

### Statistical Analysis

Statistical analysis was obtained with Student’s t-test or ANOVA (ANalysis Of VAriance) followed by Tukey’s multiple comparison’s *post-hoc* test. Analyses were performed with GraphPad Prism 5.0 or 6.0. P values are reported as n.s. (not significant) for p > 0.05, * for p < 0.05, ** p for p < 0.01, *** for p < 0.001 and **** for p < 0.0001.

## Author contributions

BBa and SS designed and performed experiments analysed the results and wrote the manuscript. BBo helped with biochemistry and discussions, CL designed the macro for FA analysis and helped with SIM images and discussions. BBo and CL read and corrected the manuscript. SEM supervised the project, interpreted the results and wrote the manuscript.

## Acknowledgements

This work was supported by La Ligue contre le cancer, the Centre National de la Recherche Scientifique and the Institut Pasteur. BB was funded by La Ligue contre le Cancer. SS is funded by the ITN PolarNet Marie Curie grant and is part of the Ecole Doctorale Frontières du Vivant (FdV) – Programme Bettencourt. We would like to thank members of the SEM lab for support and discussion, as well as JB Manneville for stimulating discussion and careful reading of the manuscript. We thank E.M. Vallés (Institut Curie), Carsten Janke (Institut Curie), Bruno Goud, (Institut Curie), and Philippe Chavrier (Institut Curie) for reagents and helpful discusions. We gratefully acknowledge JY Tinevez and A Salles and the Imagopole of Institut Pasteur (Paris, France) as well as the France-BioImaging infrastructure network supported by the French National Research Agency (ANR-10 – INSB - 04; Investments for the Future) and the Région Ile-de-France (program Domaine d’Intérêt Majeur-Malinf for the use of the Elyra microscope.

